# Comparing arm to whole-body motor control disambiguates age-related deterioration from compensation

**DOI:** 10.1101/2024.02.16.576683

**Authors:** Robin Mathieu, Florian Chambellant, Elizabeth Thomas, Charalambos Papaxanthis, Pauline Hilt, Patrick Manckoundia, France Mourey, Jérémie Gaveau

## Abstract

As the global population ages, it is crucial to understand sensorimotor compensation mechanisms. These mechanisms are thought to enable older adults to remain in good physical health, but despite important research efforts, they remain essentially chimeras. A major problem with their identification is the ambiguous interpretation of age-related alterations. Whether a change reflects deterioration or compensation is difficult to determine. Here we compared the electromyographic and kinematic patterns of different motor tasks in younger (n = 20; mean age = 23.6 years) and older adults (n = 24; mean age = 72 years). Building on the knowledge that humans take advantage of gravity effects to minimize their muscle effort, we probed the ability of younger and older adults to plan energetically efficient movement during arm-only and whole-body movements. In line with previous studies and compared to younger adults, muscle activation patterns revealed that older adults used a less efficient movement strategy during whole-body movement tasks. We found that this age-related alteration was task-specific. It did not affect arm movements, thereby supporting the hypothesis that healthy older adults maintain the ability to plan energetically efficient movements. More importantly, we found that the reduced whole-body movement efficiency was correlated with kinematic measures of balance control (i.e., the center-of-mass movement amplitude and speed). The more efficient the movement strategy, the more challenging the balance. Overall, these results suggest that reduced movement efficiency in healthy older adults does not reflect a deterioration but rather a compensation process that adapts movement strategy to the task specificities. When balance is at stake, healthy older adults prefer stability to energy efficiency.

## Introduction

Living old and healthy, also known as successful aging, is a blessing but is nonetheless associated with deterioration in various organs and functions. In terms of motor deterioration, aging is associated with loss of muscle mass (Larsson et al., 2019), sensory receptor degradation (Goble et al., 2009; Zalewski, 2015; Saftari & Kwon, 2018), and cortical atrophy (Hoffstaedter et al., 2015; Salat, 2004). Functionally, this translates into a decline in muscle strength and power (Larsson et al., 2019; Pousson et al., 2001) and movements that tend to become slower and more variable (Buckles, 1993; Darling et al., 1989). If the deteriorations are too great, they lead to reductions in quality of life and, ultimately, to dependency. Importantly, successful aging is thought to depend on compensatory processes that offset deteriorations (Baltes & Baltes, 1990; Martin et al., 2015; Zhang & Radhakrishnan, 2018). Even the most elementary concept of health includes compensatory processes at its core. The World Health Organization defined health as “a state of complete physical, mental, and social well-being and not merely the absence of disease or infirmity” (1948). Scientists and clinicians later redefined it even more generally as “the ability to adapt and to self-manage” (Huber et al., 2011; The Lancet, 2009). So, despite the normal deterioration associated with age, compensatory processes enable older adults to adapt and remain in good health (i.e., aging successfully) and thus continue to live comfortably.

In a world with a rapidly aging population (Rudnicka et al., 2020), it is essential to understand the compensatory processes that enable older people to remain healthy. This represents a critical step toward implementing interventions aimed at detecting, preventing, or reducing frailty and later dependency (for reviews, see Barulli & Stern, 2013; Ouwehand et al., 2007; Poirier et al., 2021; Zhang & Radhakrishnan, 2018). Compensation has long been theorized and could be defined as “a response to loss in means (resources) used to maintain success or desired levels of functioning (outcomes)” (Baltes, 1997). In contexts of severe deterioration, the most basic form of compensation is the use of external aids (e.g., a crutch for walking). Such compensations are observed in frail or dependent older adults, i.e., when deterioration is severe. When considering more subtle deterioration levels, identifying compensation becomes challenging. In these cases, compensatory processes enable older adults to maintain behavioral performances similar to those of younger adults, at least for the less demanding tasks of daily life (Barulli & Stern, 2013). These compensatory processes are the result of neurophysiological and behavioral adaptations that are more difficult to observe with the naked eye. Almost thirty years ago, in his famous theory of selection, optimization, and compensation, Paul Baltes and his colleagues already noted this difficulty (Baltes, 1997; Baltes & Baltes, 1990).

Since then, countless studies have explored compensatory processes using powerful tools and analyses (for recent reviews, see Bunzeck et al., 2024; Fettrow et al., 2021; Poirier et al., 2021). These studies have considerably advanced the description and understanding of age-related neural alterations. Nevertheless, behavioral compensatory processes and their underlying neural mechanisms remain essentially chimeras. Building on the theoretical work of Krakauer et al. (2017), we recently proposed that an important reason for this failure may be that studies focusing on age-related neural alterations have used overly crude behavioral paradigms (Poirier et al., 2021). Typically, these studies have used broad measures such as muscle strength, reaction time, or movement time. Although these measures and paradigms tested important functional motor performances, they measured the combination of several behavioral strategies and subtending neural mechanisms. Since these strategies and mechanisms likely showed different levels of age-related deterioration, previous studies have likely mixed deterioration and compensation processes (Poirier et al., 2021). Identifying neural compensation requires linking the brain to behavior, and to establish a precise link, we need fine behavioral measures and experimental paradigms that allow approaching the constituent processes of a behavior (Krakauer et al., 2017; Pereira et al., 2020; Urai et al., 2022). It is therefore essential to first develop detailed knowledge of age-related compensation at the behavioral level.

We sought to fill this gap by building upon the results of two different bodies of literature. On one hand, several studies have demonstrated that the brain plans efficient arm movements that take advantage of the mechanical effects of gravity to save muscle effort, thus to save energy (Berret et al., 2008; Crevecoeur et al., 2009; Gaveau et al., 2014, 2016, 2021; Gaveau & Papaxanthis, 2011; Gueugneau et al., 2023; for a review, see White et al., 2020). Importantly, recent work demonstrates that this ability is maintained and maybe even upregulated in older adults (Healy et al., 2023; Huang & Ahmed, 2014; Poirier et al., 2020; Summerside et al., 2024). On the other hand, studies probing the control of movements performed with the entire body have reported a different conclusion. Kinematic results suggest that older adults plan whole-body movements that are less energy-efficient than younger adults (Casteran et al., 2018; Paizis et al., 2008). This is unexpected because such movements require more energy expense in older adults than in younger adults (Hortobagyi et al., 2003, 2011; Julius et al., 2012; VanSwearingen & Studenski, 2014). Since the ability to plan efficient movements is maintained in older adults, as testified by arm movements studies, one may speculate that this decreased efficiency reflects an age-related compensation that changes movement strategy (i.e., an age-related motor adaptation process). However, because this literature used very different experimental paradigms and measurements, this conclusion is highly speculative. More importantly, the results of numerous other studies could also interpret the decreased efficiency observed in whole-body movements as a deterioration of the ability to produce efficient motor patterns (Goodpaster et al., 2006; Henry & Baudry, 2019; Quinlan et al., 2018; Vernazza-Martin et al., 2008). Here we test the hypothesis that age-related alterations in movement efficiency correspond to an adaptation process, i.e., a change in movement strategy that compensates for other deteriorated sensorimotor components. To overcome the aforementioned limitations, using a specific muscle activation pattern analysis that has proven relevant to focusing on this precise process of energetic efficiency (Chambellant et al., 2023; Gaveau et al., 2021; Poirier et al., 2022, 2023a; Thomas et al., 2023), we compare older to younger adults on tasks involving either arm or whole-body movements. We then test whether energy efficiency is correlated to balance control.

## Methods

### Participants

Because we had no prior data to calculate the ideal sample size, we included as many participants as possible over a fixed recruitment period. Twenty younger adults (23.6 ± 2.1 y.o.) and twenty-four older adults (72 ± 5.3 y.o.) were included in the study after giving their oral informed consent. Participants had normal or corrected-to-normal vision and did not present any neurological or muscular disorders. The laterality index of each participant was superior to 60 (Edinburgh Handedness Inventory, Oldfield 1971), indicating that all participants were right-handed. The study was carried out following legal requirements and international norms (Declaration of Helsinki, 1964) and approved by the French National Ethics Committee (2019-A01558-49). Each participant was included in the study by a medical doctor.

### Experimental Protocol

All participants performed four tasks in a randomized order. These tasks either required moving the arm only (Figure 1A) or the whole-body (Figure 1B-D). Whole-body movements consisted of seat-to-stand/back-to-sit (STS/BTS, Figure 1B), whole-body reaching toward near targets (WBR D1, Figure 1C), and whole-body reaching toward far targets (WBR D2, Figure 1D). The arm task was selected because it is the reference task that has been studied to demonstrate how muscle patterns take advantage of gravity effects to save energy. The whole-body tasks were selected because they include an equilibrium constraint, are movements of the daily life, and they have been investigated in previous studies (Casteran et al., 2018; Jeon et al., 2021; Manckoundia et al., 2006; Millington et al., 1992; Mourey et al., 1998; Paizis et al., 2008).

**Figure 1.**
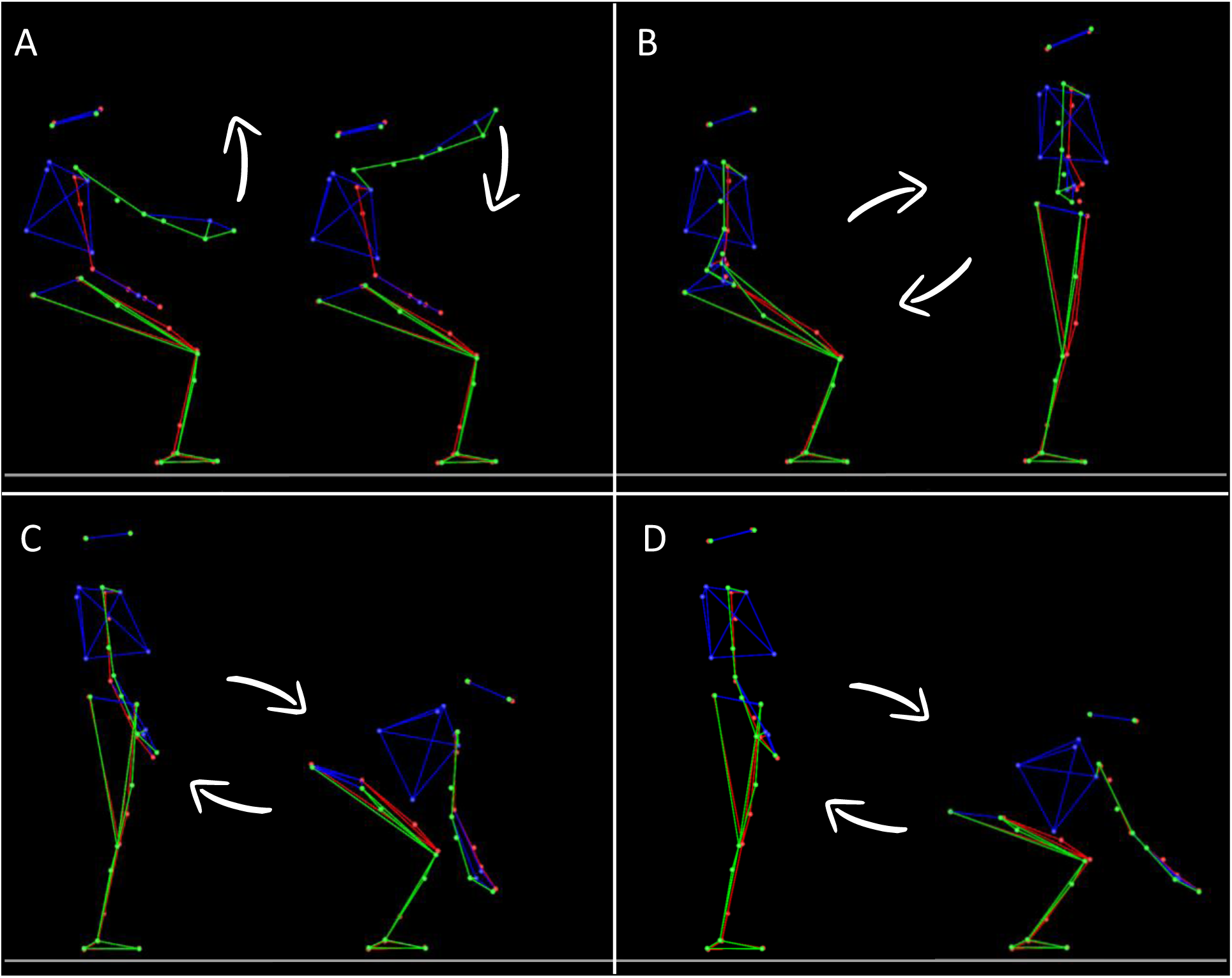
Illustration of the four tasks. Each panel illustrates the extreme body positions between which participants performed their movements. Each position was alternatively the starting or ending target of a movement, depending on movement direction. A: Single degree of freedom arm movements flexion/extension around the shoulder joint (flexion/extension). Participants performed upward and downward arm movements. B: Seat-to-stand/Back-to-sit movements. Participants performed vertical multi-articular whole-body movements to either stand up from the stool (upward movement) or sit on it (downward movement). C: Whole-body reaching task toward a near target. Participants performed vertical multi-articular whole-body movements to either reach towards targets that were located nearby the floor (downward movement) or to bounce back from this position toward a resting vertical standing position (upward movement). D: same as C but with targets that were placed farther away on the antero-posterior axis.

#### ARM task

Over a variety of arm movement tasks, including single or muti-degree of freedom pointing movements, drawing movement, reach to grasp movements, or arm movements that transport a hand-grasped object, the results consensually support an optimization principle that shapes arm motor patterns to take advantage of gravity effects in saving energy (Crevecoeur et al., 2009; Gaveau et al., 2011; Gaveau & Papaxanthis, 2011; Le Seac’h & McIntyre, 2007; Paizis et al., 2008; Papaxanthis et al., 1998, 2005; Yamamoto & Kushiro, 2014a). Thus, to make the protocol doable in a single session with each participant, we only included one arm task in the present experiment. This task was similar to a task used in several previous studies probing human movement adaptation to the gravity environment (Gaveau et al., 2014, 2016, 2021; Gaveau & Papaxanthis, 2011; Gentili et al., 2007; Hondzinski et al., 2016; Le Seac’h & McIntyre, 2007; Poirier et al., 2020, 2022; Yamamoto & Kushiro, 2014a). Using their right arm, participants carried out single-degree-of-freedom vertical arm flexion/extension movements around the shoulder joint. Two blocks of arm movements were performed in a randomized order. One block consisted of six slow movements, and one block consisted of twelve fast movements. Two targets (diameter of 3 cm) were placed in front of the participant’s right shoulder (in a parasagittal plane) at a distance corresponding to the length of their fully extended arm plus two centimeters. The prescribed movement amplitude between the two targets was 45°, corresponding to 112.5° (upward target, 22.5° above horizontal) and 67.5° (downward target, 22.5° below horizontal) shoulder flexion/extension.

#### STS/BTS task

This task was similar to those of previous studies (Jeon et al., 2021; Manckoundia et al., 2006; Millington et al., 1992; Mourey et al., 1998). Participants were seated on an armless stool whose height was adjusted to correspond to 30% of the participant’s height. The hands were positioned on the hips, and the back was instructed to be maintained about vertical. Participants were asked to stand up from the stool, make a short pause (about 2s), and then sit back on the stool. Similarly to arm movements, participants executed two blocks of movements in a randomized order. One block consisted of six slow movements, and the other consisted of 12 fast movements.

#### WBR task

This task was similar to those of Casteran et al. (2018) and Paizis et al. (2008). Starting from an upright position, we asked participants to perform whole body reaching movements (WBR) toward two targets nearby the floor with their two index fingers (10% of their heights above the floor). The two targets (4 × 2 cm) were spaced by 0.5 m on a medio-lateral axis and centered on the participant’s sagittal axis. They were placed in front of the participant at two different distances, corresponding to 15% (D1) or 30% (D2) of their height on the antero-posterior axis. Distances were measured from the participant’s big toe. Similarly to the previous two tasks, for each distance and in a randomized order, participants executed two blocks of trials performed at two different speeds. One block consisted of six slow movements and the other twelve fast movements (total of four blocks: two speeds × two distances).

#### Trial organization

The organization of a trial was similar for all tasks. It was carried out as follows: i) the experimenter indicated to get ready; ii) the participant adopted the requested initial position; iii) after a brief delay (∼1 second), the experimenter verbally informed the participant that she/he was free to reach the requested final position whenever she or he wanted. Note that reaction time was not emphasized in our experiment; iv) the participant was requested to maintain the final position for a brief period (about 1 second); v) the experimenter instructed to move back to the starting position (reversed movement) whenever desired; vi) lastly, the participant was asked to relax. A short rest period (∼20 s) separated trials to prevent muscle fatigue. Additionally, participants were free to rest as long as they wanted between blocks. Participants were allowed to perform a few practice trials (∼3 trials) before each block. Low-speed and high-speed blocks were similar except that the instructions were to perform the movements in roughly 5 seconds or as fast as possible, respectively.

### Data Collection

#### Kinematics

We used the Plug-In Gait full body model (Vicon, Oxford Metrics, UK) following their recommendations to place the 39 reflective markers on the participant’s head (temples and backs of the head to form a rigid plan with the head), back (C7, T10 and on the right scapula), torso (jugular notch where the clavicles meet the sternum and on the xiphoid of the sternum), shoulders (acromion), arms (upper lateral 1/3 for the left arm, and 2/3 for the right arm), elbows (lateral epicondyle), forearms (lower lateral 1/3 for the left forearm, and 2/3 for the right forearm), wrists (both cubitus styloid processes), hands (middle of the proximal knuckle of the index), pelvis (anterior and posterior superior iliac spine), thighs (upper lateral 1/3 for the left leg, and 2/3 for the right leg), knees (lateral side of the flexion-extension axis), calves (upper lateral 1/3 for the left calf, and 2/3 for the right calf), ankles (lateral malleolus), and feet (second metatarsal head and heel). The markers on the scapula, on the arms, on the forearms, on the thighs, and on the calves have been deliberately placed asymmetrically so that the model can best dissociate the right and left sides; these markers are not used for the analyses presented in this manuscript.

We recorded the position of all markers with an optoelectronic motion capture system (Vicon system, Oxford Metrics, UK; 18 cameras) at a sampling frequency of 200 Hz. The spatial variable error of the system was less than 0.5 mm.

#### EMG

We placed sixteen bipolar surface electrodes (Cosmed, pico EMG, sampling frequency: 1000Hz) on the anterior (AD) and posterior (PD) heads of the deltoid, vastus lateralis (VL), biceps femoris (BF), spinal erectors on L1 (ESL1) and on T7 (EST7), the soleus (SOL), and on the tibialis anterior (TA) to record EMG activity. Electrodes were placed bilaterally. The location of each electrode was determined following the recommendations from Barbero et al. (2012).

The Giganet unit (Vicon, Oxford Metrics, UK) synchronously recorded kinematic and EMG data.

### Data Analysis

We processed kinematic and EMG data using custom programs written in Matlab (Mathworks, Natick, MA). Data processing was inspired by previous studies (Gaveau et al., 2021; Poirier et al., 2022) and was similar for all tasks.

#### Kinematics analysis

First, we filtered position using a third-order low-pass Butterworth filter (5 Hz cut-off, zerophase distortion, “butter” and “filtfilt” functions). We then computed the amplitude of the movement using steady phases (200ms for fast movements and 500ms for slow movements) before and after the movement, using the marker of the right shoulder (for whole-body movements, see Figure 2) or the right finger (for arm movements). The amplitude was computed on the Z axis for fast movements and on X, Y, and Z axes for slow movements. For slow movements, we used 3D position to minimize detection error on signals that were more variable than those obtained during fast movements. Last, we automatically defined movement onset and offset as the moments when the displacement rose above or felt below a threshold corresponding to 5% and 95% of the total movement amplitude, respectively.

**Figure 2.**
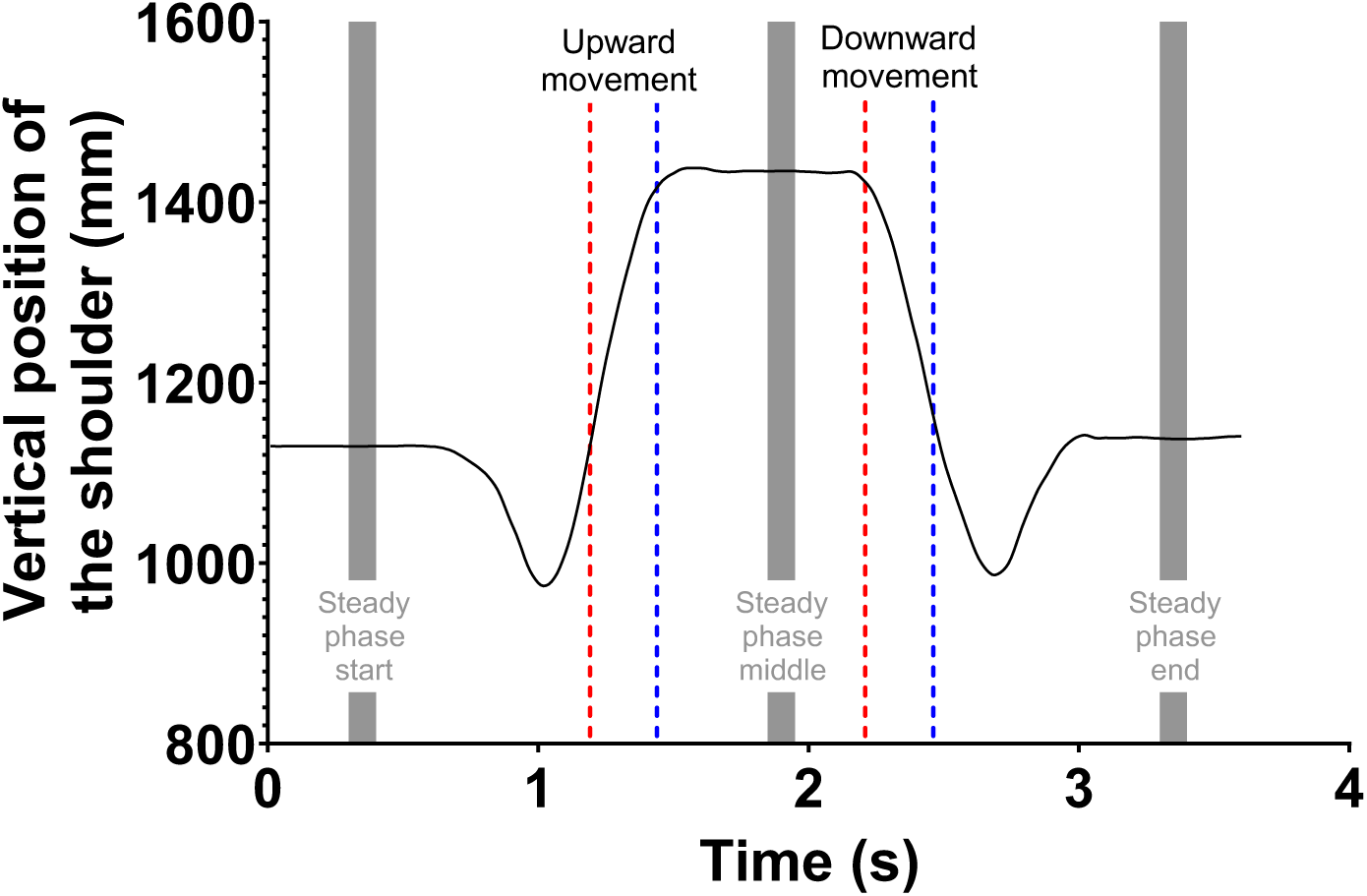
Detection of movement start and end. This panel displays the recording of two successively opposite fast Sit to stand / Back to sit movements. The black trace represents the position of a shoulder marker through time. Rest position is collected during steady phases, before and after each movement (ascending or descending). Based on the data obtained in steady phases, a recursive algorithm automatically defined movement onset and offset as the moments when the displacement rose above or felt below a threshold corresponding to 5% and 95% of the total movement amplitude, respectively.

On behalf of using the kinematics to define the start and end of movement, we analyzed the displacement of the Center of Mass (CoM) in three dimensions to understand how equilibrium was maintained during the whole-body tasks. This was done to reproduce the work of Casteran et al., (2018) and Paizis et al., (2008), but also, and more importantly, to perform a simple analysis testing whether our main criterion, quantified via electromyographic activity, is linked to a simple, interpretable kinematic change. Our analysis utilized a seven-segment mathematical model incorporating rigid segments such as the Trunk, Thigh, Shank, Foot, Upper arm, Forearm, and Hand. We used anthropometric data from Winter (2009), as performed by previous studies (Berret et al., 2009; Stapley et al., 1999). Our choice of movement segmentation for this specific kinematic analysis has been guided by the works of Casteran et al., (2018) and Paizis et al., (2008). We determined movement onset and offset on velocity profiles, using a threshold of 5% of the peak velocity. We further explored the kinematics of the whole-body tasks using two simple parameters: i) the total displacement of the center of mass, calculated as the distance between the start and end positions and normalized by the subject’s height; and ii) the peak velocity of the center of mass. We focused on downard movements, as these are the ones that have been studied and present the greatest challenge to balance. The specific process to compute criteria used by previous studies (Casteran et al., 2018 and Paizis et al., 2008) is detailed and available in Supplementary Figure 1.

#### EMG analysis

Below, following methodologies developed by several previous studies, we detail how we obtain EMG marker.

##### Pre-processing

EMG signals were first rectified and filtered using a bandpass third-order Butterworth filter (bandpass 30-300 Hz, zero-phase distortion, “butter” and “filtfilt” functions) followed by a low-pass third-order Butterworth filter (low-pass frequency: 5 Hz) to highlight important features of muscular activities. Signals were integrated using a 100ms sliding window using trapezoidal numerical integration from Matlab (Mathworks, Natick, MA) and cut off. For fast movements, EMG signals were cut off from 240ms before movement onset to 90ms after movement offset. For slow movements, EMG signals cut off from 75ms before movement onset to 75ms after movement offset. These timing values were obtained from preliminary analyses detecting EMG activity start and stop before and after all movements. The result is the average of all participants. Importantly, those values were kept constant for all participants and, thus, should not bias group comparisons.

##### Phasic/tonic separation

We then computed the phasic component of each EMG signal using a well-known subtraction procedure that has mostly been used to study arm movements (Buneo et al., 1994; d’Avella et al., 2006, 2008; Flanders et al., 1994; Flanders & Herrmann, 1992; Gaveau et al., 2021). This processing allows quantifying how much the central nervous system takes advantage of the gravity torque when moving the body in the gravity environment (Gaveau et al., 2021; Poirier et al., 2022, 2023a). Here, we customized this procedure to investigate whole body movements since movements are not one-degree-of-freedom movements. First, the tonic signal was obtained from the six slow movements. For that purpose, the cut movements (as described earlier with delays) were normalized in duration to be finally averaged together in one tonic signal. Second, to improve signal to noise ratio, EMG traces of fast movements were ordered according to movement mean velocity and averaged across two trials (from the two slowest to the two fastest movements). This resulted in six EMG traces to be analyzed for each block. Each set of two traces was normalized in duration (corresponding to the mean duration of the two traces) before averaging. Third, the phasic component was obtained by subtracting the tonic EMG from the EMG trace of each pair of fast movements. Finally, to set the data of all participants on a common scale, phasic activity was normalized by the maximal EMG value recorded in each task for each participant.

##### Muscles selection

It was recently shown that the phasic EMG activity of antigravity muscles, those that pull against the gravity vector, consistently exhibits negative epochs (Chambellant et al., 2024; Gaveau et al., 2021; Poirier et al., 2022; Thomas et al., 2023) when the arm acceleration sign is coherent with the gravity acceleration sign (i.e., in the acceleration phase of downward movement and in the deceleration phase of upward movements). This observation likely reflects an optimal predictive motor strategy where muscle activity is decreased when gravity assists arm movements, thereby saving energy (Gaveau et al., 2021). In the present study, the antigravity muscles are: i) the Anterior Deltoïd (DA), flexing the shoulder joint; ii) the Vastus Lateralis (VL), extending the knee joint; iii) the Erector Spinae L1 (ESL1), extending the rachis; iv) the Erector Spinae T7 (EST7), extending the rachis; v) the Soleus (SOL), flexing the ankle in the plantar direction. Because the Erector Spinae T7 and the Soleus muscles did not play a strong focal role but a rather postural one in the present tasks, we focused our analyses on the remaining three muscles (DA, VL, and ESL1). Probing the activation of a postural muscle, per definition, is not appropriate to test whether the nervous system takes advantage of gravity effects to move our body limbs. Compared to other joints (e.g., hips and knees), the ankle and upper rachis were only minimally mobilized in the tasks we investigated here (see stick diagrams in Figure 1). Including these muscles in our analyses would thus add noise to our dependent variables and likely impede our ability to test our hypothesis. Therefore, we focused on DA during arm movements and on VL and ESL1 during movement of the entire body.

##### Quantifying negativity

We defined negative epochs as an interval where the phasic EMG signal was inferior to zero minus three times the standard deviation of the stable phase preceding the movement, and this for at least 40ms. This duration has been chosen after preliminary tests to avoid detecting false-positives. We kept it constant for all analyses. We used this value as a threshold to automatically detect negativity onset and offset. On each negativity phase, we computed: i) a negativity index, defined as T x NA / TA, with NA the Negative Area integrated on the phasic signal between negativity onset and offset, TA the Tonic Area integrated on the tonic signal between the negativity onset and offset, and T the duration of the negative epoch normalized by movement duration (see Figure 3). This value is always negative or null. The lower the value, the greater the efficiency; ii) negativity occurrence, defined as the number of trials where a negative epoch was automatically detected, divided by the total number of trials in the condition; iii) negativity duration, defined as the duration of the negative epoch normalized by movement duration; iv) negativity amplitude, defined as the minimal *Phasic value / Tonic value* × 100 during the negative period. A value of −100 indicates that the muscle is completely relaxed and a value of 0 indicates that the muscle exactly compensated the gravity torque.

**Figure 3:**
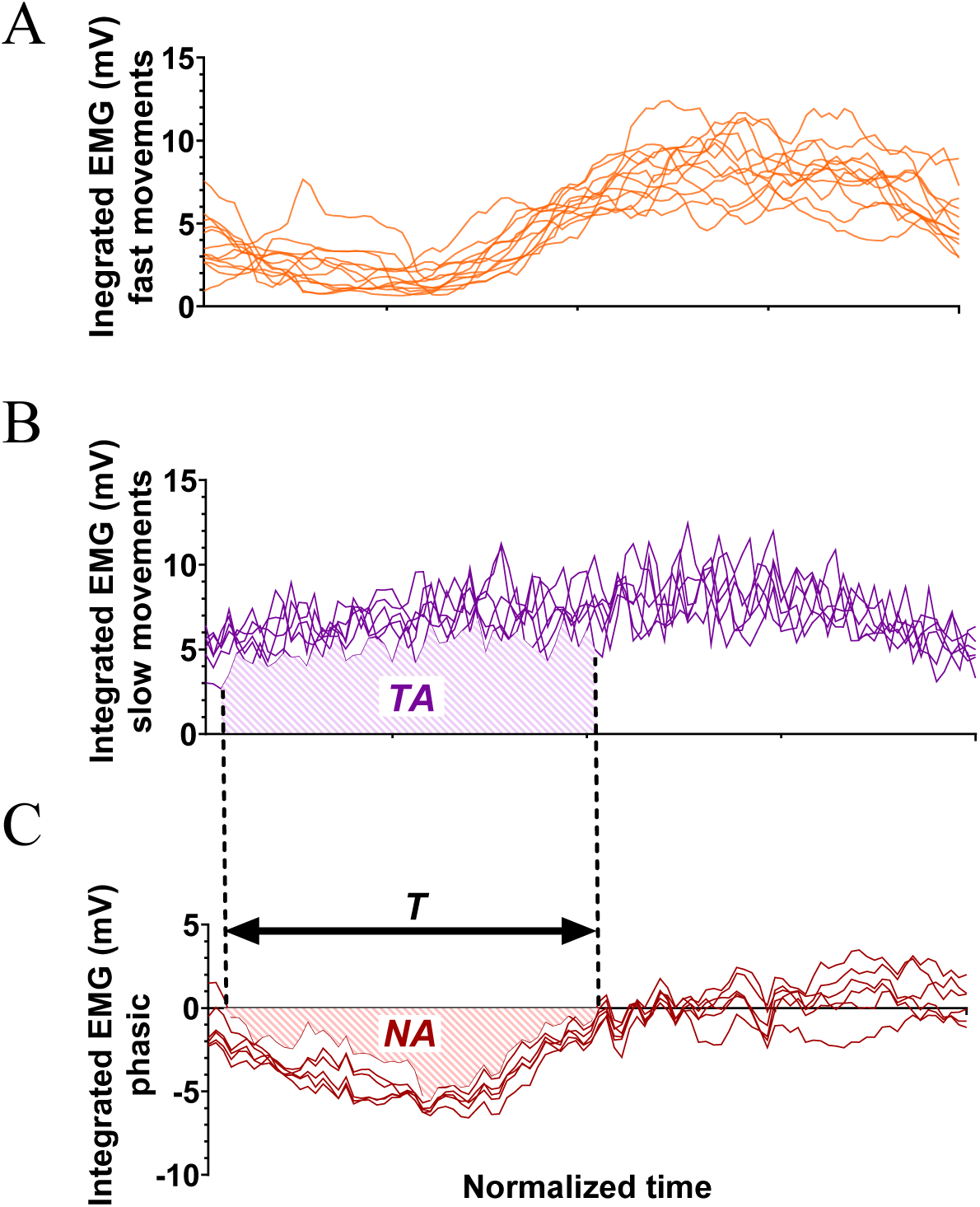
Illustration of the calculation method to obtain phasic EMG components. Electromyographic signals (mV) are presented as a function of time. Pattern duration and amplitude are normalized (see methods). **A**: Six integrated Vastus Lateralis EMG signals during fast BTS movements of a typical participant (BTS: Back-to-seat); **B**: Six integrated Vastus Lateralis EMG signals recorded during slow BTS movements of a typical participant. These signals represent the tonic component. TA: Tonic Area integrated on the tonic signal between the negativity onset and offset; **C**: Integrated phasic EMG component computed using the six fast (panel **A**) and slow movements (panel **B**). The phasic is calculated by subtracting the mean of the slow acquisitions from the fast acquisitions (Phasic = Fast – Tonic). T: the duration of the negative epoch normalized by movement duration and NA: the Negative Area integrated on the phasic signal between negativity onset and offset.

As is often the case with EMG recordings, some of the EMG signals exhibited aberrant values. Those signals are usually due to poor contact between the electrodes and the skin. Supplementary Table 1 summarizes the issues encountered with all electrodes and participants.

### Machine Learning

We used custom Matlab (Mathworks, Natick, MA) scripts to perform all machine learning analyses. The ESL1G was not considered for these analyses because the electrode was defective for several younger participants (see Supplementary Table 1).

The input data was the phasic EMG signals of the 15 muscles taken individually or the whole set at once. These vectors were fed to the algorithms using binary classification setups, where the algorithm learned to distinguish between the EMGs of the two groups. To ensure robustness of the results, we employed a five-fold cross-validation method. This involved splitting the whole dataset into five sets while ensuring equal representation of both directions in each set. The algorithm was trained on four of those sets before being evaluated on the fifth set (containing data unknown to the trained algorithm). This operation was repeated five times, so each set was tested once. Cross-validation allowed computing the average accuracy and its variance across the testing sets, thereby providing a reliable estimate of the accuracy obtained by the algorithm. Finally, we could compare the accuracy of the algorithm for each muscle.

### Univariate Statistics

After an initial kinematic analysis (detailed in the results section), we observed a difference in movement duration between younger and older adults (conducting a repeated measure analyses of variance with a between factor *Age* with two levels: Young/Older and a within factor *Task-type* with two levels: Arm/Whole-body movements). Because movement duration is known to influence phasic EMG negativity (Poirier et al., 2023b), we added movement duration as a covariate. We performed repeated measure analyses of covariance (ANCOVA) using JASP software. Two ANCOVA analyses were carried out. We first used a mixed ANCOVA with a between factor *Age* (two levels: Young/Older) and a within factor *Task-type* (two levels: Arm/Whole-body movements) to test whether age effects on movement control depended on the type of task being performed. Second, to detail the age differences observed during movements of the entire body, we used a mixed ANCOVA with a between factor *Age* (two levels: Young/Older) and a within factor *Whole-Body-Tasks* (three levels: STS_BTS/WBR D1/WBR D2). In all cases, the significance level was set to 0.05.

To test for a possible beneficial effect (i.e., compensation) of the EMG alterations that we observed with aging, we performed a kinematic analysis of the center of mass. We then used independent Student-tests and Pearson correlation coefficients to study potential differences between groups and associations between variables.

## Results

Movement duration of fast movements varied between tasks and was slightly reduced in younger compared to older participants (see Figure 4 and Supplementary Table 2 for detailed results). Overall, older adults were 3.5% slower than younger adults. A repeated measures ANOVA revealed that this age-difference was significant (F_(1,42)_= 14.5, P=4.58E-05, ƞ^2^=0.256). For this reason, we used movement duration as a covariate in the following statistical analyses. Nevertheless, as revealed by Figure 4, it is important to note that an important number of older adults moved with durations that were similar to those of younger adults.

**Figure 4.**
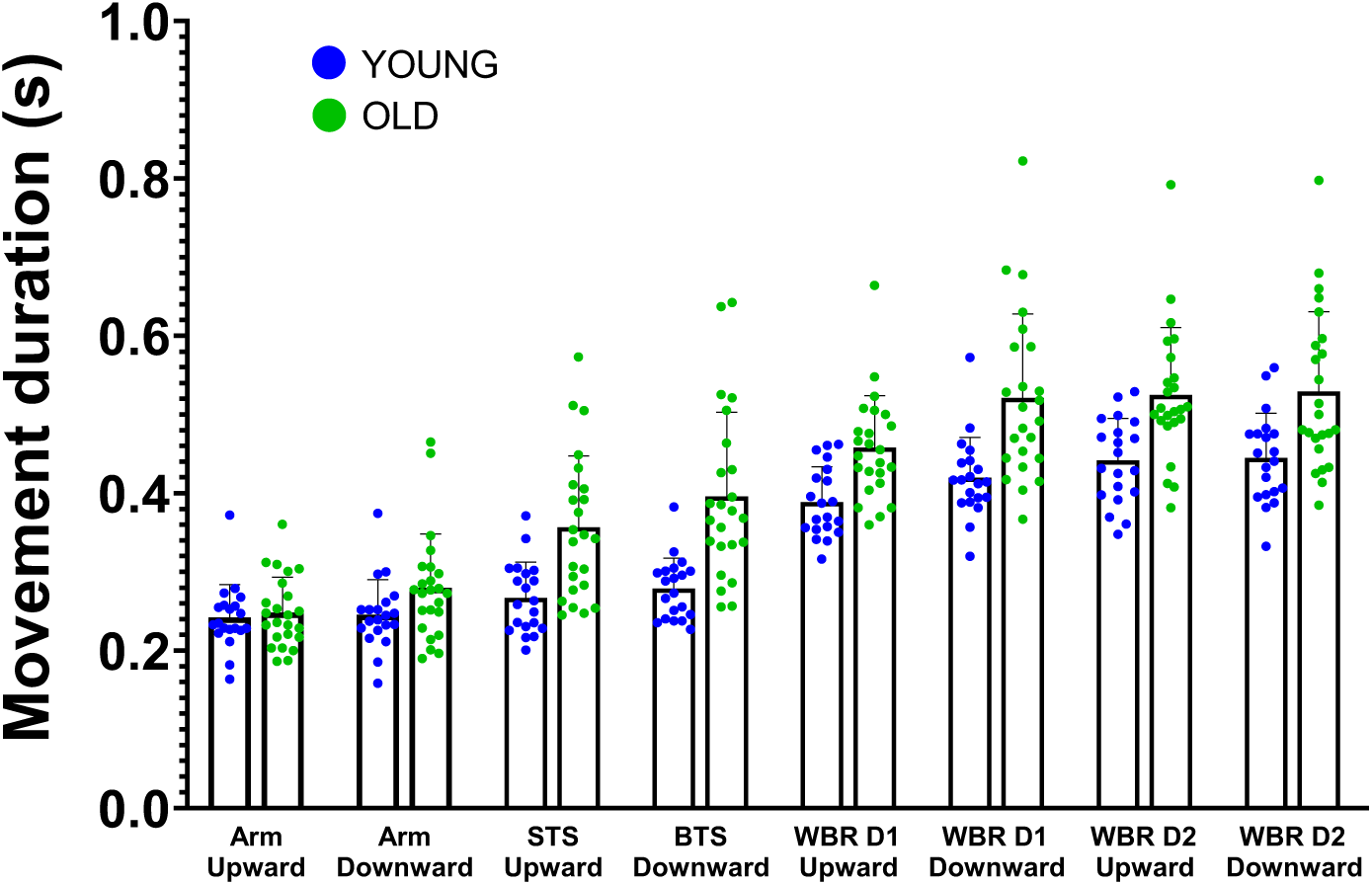
Mean ± SD movement durations (s) for fast movements performed in all tasks and both groups (STS: Seat-to-stand, BTS: Back-to-seat, WBR: Whole-body-reaching, D1: Short distance=15% of the height of the subject, and D2: Long distance=30% of the height of the subject). Each point corresponds to the average duration of the trials of one participant. The blue points represent the young participants, and the green points correspond to the older participants.

A body of computational studies has demonstrated that human arm movements take advantage of gravity effects to save energy (Berret et al., 2008; Crevecoeur et al., 2009; Gaveau et al., 2011, 2014, 2016, 2021). Most of these studies used mathematical models that minimized the absolute work of muscle force to produce the arm displacement (Berret et al., 2008; Gaveau et al., 2011, 2014, 2016, 2021). More specifically, the study from Berret et al. (2008) formally demonstrated that this muscle work cost, and associated behavior, corresponded to an energetic-like optimum. Because previous studies have shown that the amplitude of kinematic and electromygographic markers directly relates to energetic efficiency (Gaveau et al., 2016; Poirier et al., 2022), here we compare the amplitude of an EMG marker between younger and older adults. If the EMG marker increases, this means that energetic efficiency increases – i.e., the minimization process is upregulated – and thus muscle work decreases. If the EMG marker decreases, this means that energetic efficiency decreases – i.e., the minimization process is downregulated – and thus muscle work increases.

Figure 5 displays average phasic EMG profiles for each muscle, direction, and task. As recently reported, phasic EMG signals of arm movements show negative phases during the deceleration of upward and the acceleration of downward arm movements, i.e., where gravity torque helps generate the arm’s motion (Gaveau et al., 2021; Poirier et al., 2022, 2023a). Previous works demonstrated that this negativity is not erratic but systematic and indicate that muscles contract less than necessary to compensate for gravity effects. It is therefore especially prominent on antigravity muscles and reveals that the central nervous system (CNS) exploits gravity effects to produce efficient movements, i.e., motor patterns that save unnecessary muscle work. Here, we extend this result to movements performed with the entire body. Indeed, for STS/BTS and WBR movements, Figure 5B-D unveils phasic EMG negativity during the deceleration of upward movements and the acceleration of downward movements, i.e., when gravity can help produce the motion. This first qualitative result demonstrates that movements that are performed with the entire body, similarly to more focal arm movements, exploit gravity effects to save unnecessary muscle work (Gaveau et al., 2021). More importantly, the present results qualitatively reveal that older adults also use such an efficient strategy, both when moving their arm and their entire body.

**Figure 5.**
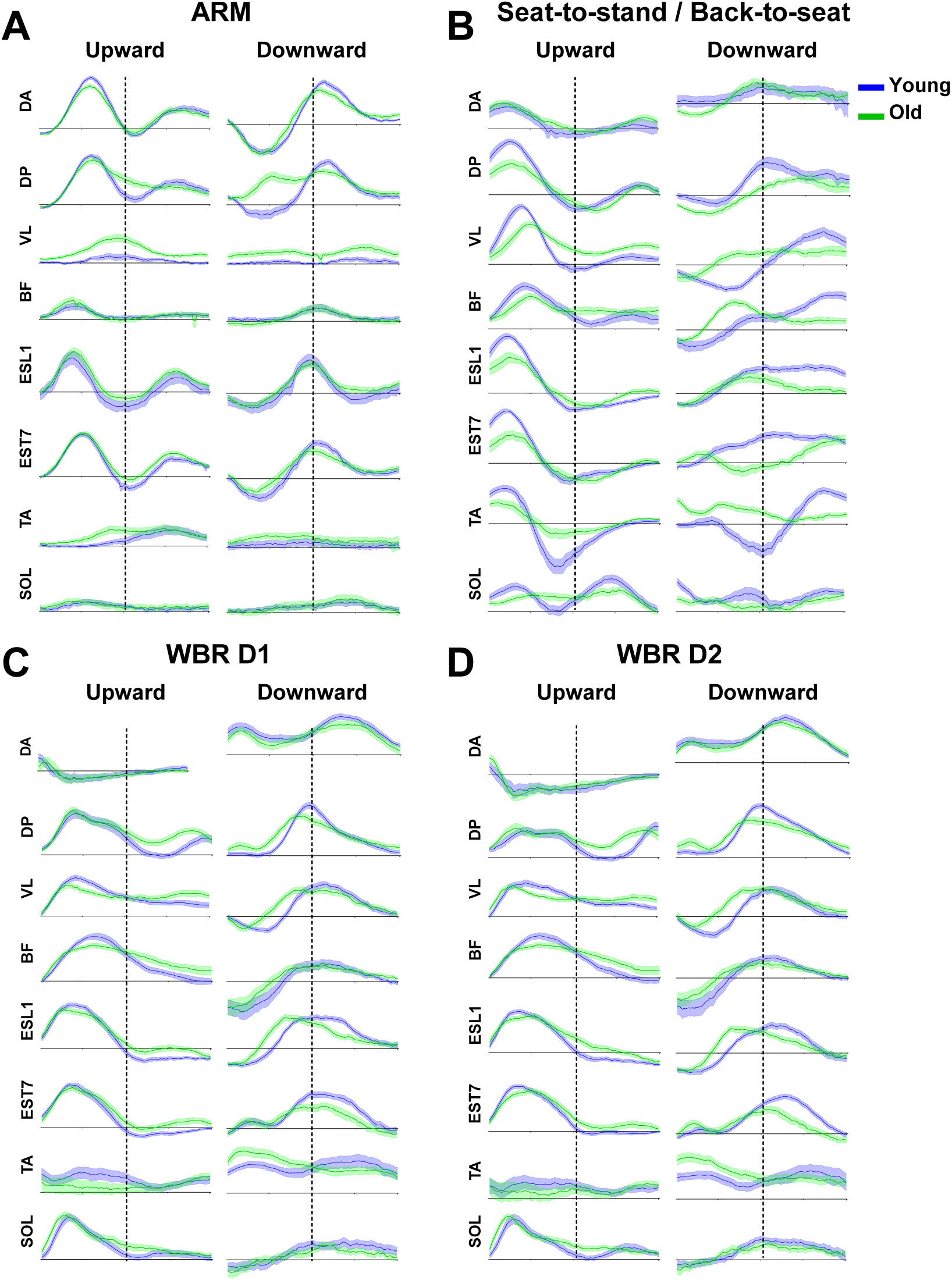
Mean (±SE) integrated phasic EMGs recorded for both groups (n = 20 for younger and n = 24 for older) during arm (panel A), Seat-to-stand/Back-to-seat (panel B), and whole body reaching movements (panel C for short distance D1 and panel D for long distance D2). Blue traces present EMGs recorded for younger participants, while green traces present EMGs recorded for older participants. The dotted line divides the movement in two: the first half is acceleration and the second is deceleration. (DA: Anterior deltoid, DP: Posterior deltoid, VL: Vastus Lateralis, BF: Biceps Femoris, ESL1: Erector Spinae in L1, EST7: Erector Spinae in T7, TA: Tibialis Anterior, SOL: Soleus).

### Main analysis

Following our primary hypothesis, we first analyzed a single metric quantifying phasic EMG negativity on an average muscle activation pattern (vastus lateralis and erector spinae in L1 were averaged for whole-body tasks and deltoid anterior was used for arm tasks), namely the negative area of phasic EMG patterns (see methods and Poirier et al., 2022, 2023a). The bigger the negativity index, the more efficient the muscle contractions, in the sense that gravity effects were maximally exploited to save energy (Gaveau et al., 2021). Figure 6 displays the results of this ANCOVA analysis (Age *×* Task-Type), revealing a significant interaction between age and task factors (F_(1,42)_= 5.48, P=2.44E-02, ƞ^2^=0.120) but no Age or Tasks effect (for detailed statistical results, please see Supplementary Table 3). This result demonstrates that age differently alters motor strategies for arm movements vs whole-body movements. Older adults used gravity effects to a similar extent as younger ones when performing arm movements (older adults, mean ± SD: −10.7 ± 5.6, 95% CI: [-8.4;-13.0]; younger adults, −11.4 ± 3.6, [-9.8;-13.0]), but to a lesser extent when performing whole-body movements (older adults, −9.7 ± 3.2, [-8.0;-11.5]; younger adults, −15.6 ± 3.3, [−14.1;−17.0]). As recently reported by Poirier et al. (2023a), similar arm results in younger and older adults suggest that the ability to plan movements that optimally use gravity effects to save energy remains functional in older adults. The results obtained in whole-body movement tasks (STS/BTS and WBR) could thus suggest that the difference observed between older and younger adults does not reflect a deterioration of the ability to plan movements that are optimally adapted to the gravity environment. Instead, it would suggest a change in movement strategy that compensates for other deteriorated control processes (for example, the loss of muscle mass & force).

**Figure 6.**
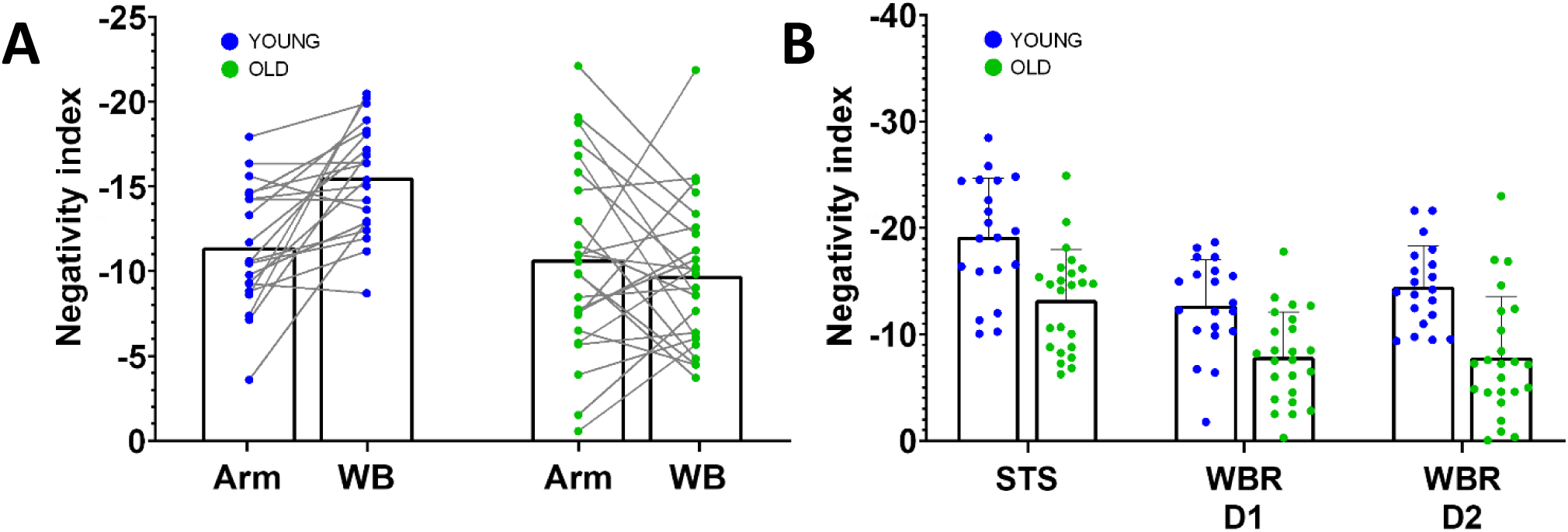
Negativity index. Computed for **A**. arm and whole-body movements in both groups (WB: Whole Body combines seat-to-stand/back-to-seat, whole-body reaching from D1 and whole-body reaching from D2) and **B**. each whole-body task (STS/BTS: seat-to-stand/back-to-sit, WBR D1: whole-body reaching from D1 and WBR D2: whole-body reaching from D2). The negativity index, defined as T x NA / TA, with NA the Negative Area integrated on the phasic signal between negativity onset and offset, TA the Tonic Area integrated on the tonic signal between negativity onset and offset, and T the duration of the negative epoch normalized by movement duration. The blue points correspond to the younger participants, and the green points correspond to the older participants. Each point corresponds to the mean value of one participant (mean across trials and antigravity muscles, and/or tasks).

We performed a complementary analysis to determine whether every whole-body task showed the same age effect (ANCOVA Age x Whole-Body Tasks). This test did not reveal any interaction effect (F_(2,42)_= 0.77, P=4.67E-01, ƞ^2^=0.019), further supporting the interpretation that this is the “whole body” aspect of the task that impacts the motor strategy in older adults (please see Supplementary Table 3 for full analysis).

Previous studies have proposed that the change in kinematic strategies observed between older and younger adults during whole-body movements could be explained as a strategy maximizing equilibrium maintenance rather than energetic efficiency (Casteran et al., 2018; Paizis et al., 2008). Following this hypothesis, one would predict increasing differences between younger and older adults when the equilibrium constraint increases. In the present experiment, increased equilibrium constraint was produced by increasing the target distance during whole body reaching movements (WBR D1 vs WBR D2; alike Casteran et al., 2018). The Age x Whole-Body Tasks ANCOVA, however, did not reveal such a difference.

Last, we analyzed kinematic patterns in order to investigate whether the decreased energetic efficiency observed during whole body tasks in older adults could actually be interpreted as compensation. We tested whether the negativity of phasic EMGs correlated with kinematic parameters that are related to balance control (the COM displacement, and COM peak velocity, see Figure 7; and see Supplementary Figure 1 for detailed results of the reproduction of the tests conducted by Casteran et al., (2018) Paizis et al., (2008). The EMG criterion during the Back to Seat task was found to be significantly correlated with the COM displacement (Pearson correlation, P=2.2E-2, Pearson^′^s r=-0.343) and the COM peak velocity (Pearson correlation, P=1.9E-3, Pearson^′^s r=-0.476). This same EMG criterion also turned out to be significantly correlated for the Whole-Body Bending task with the COM displacement (Pearson correlation, P=3.2E-3, Pearson^′^s r=-0.435) and with the COM peak velocity (Pearson correlation, P=1.2E-7, Pearson^′^s r=-0.700). The linear regressions revealed that the more a participant used the effects of gravity, the more and the quicker she/he displaced his COM. One could interpret this result as demonstrating that older adults lose the ability to plan energetically efficient movement and, thus, move their whole-body less and more slowly. However, the null age effect on arm movement control supports the hypothesis that planning efficient movements remains functional in older adults, as also supported by recent other results (Healy et al., 2023; Huang & Ahmed, 2014; Poirier et al., 2020; Summerside et al., 2024). Overall, during movements performed with the entire body, i.e., when equilibrium maintenance is challenged, the present results support an age-related adaptation process that selects a less energetically efficient but more stable movement strategy in healthy older adults.

**Figure 7.**
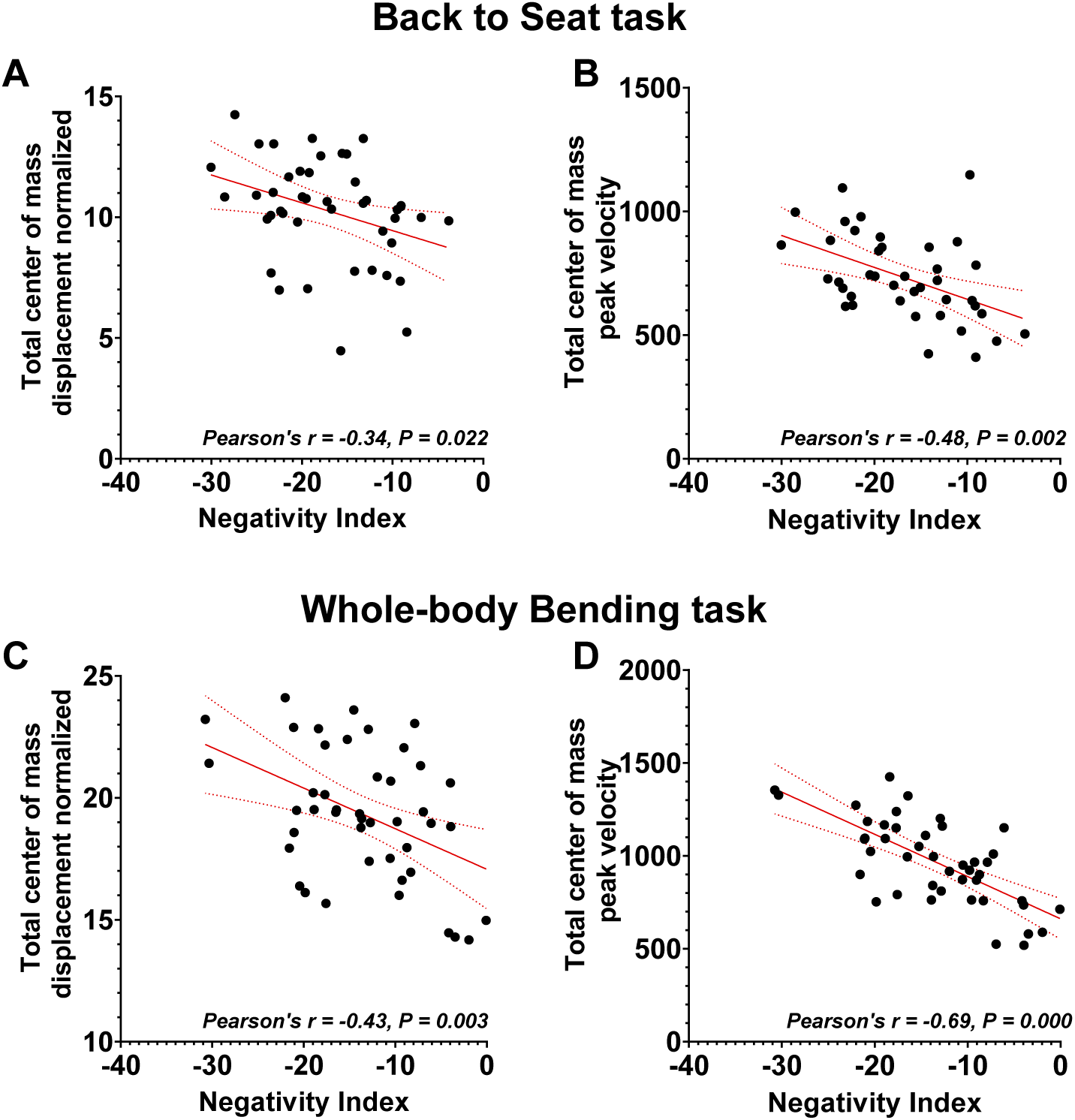
Center of mass analyses. Linear relationship between EMG negativity (Vastus Lateralis and Spinal Erector L1) and total displacement of the center of mass during A. back-to-sit movements and B. bending movements from the whole-body reaching tasks (averaged between distances D1 and D2).

### Exploratory analyses

To provide a fine-grained analysis of the age effect on phasic EMG negativity during whole-body motion, we probed negativity duration, negativity amplitude, and negativity occurrence across tasks and age-groups (see Figure 9). Here also, the bigger the values, the bigger the use of gravity effects to produce body motion. A repeated measure ANCOVA Age x Tasks (Young/Older x STS_BTS/WBRD1/WBRD2) revealed a significant age effect where negativity duration was larger in younger compared to older participants (F_(1,36)_= 21.49, P=4.54E-05, ƞ^2^=0.374). The age effect did not reach significance for negativity occurrence (F_(1,36)_= 3.62, P=0.065, ƞ^2^=0.091) nor for negativity amplitude (F_(1,36)_= 1.16, P=0.28, ƞ^2^=0.031). No interaction between Age and the other factors reached significance (see Supplementary Table 3 for detailed results). Overall, all variables showed qualitatively smaller negativity on phasic EMGs, thus reduced use of gravity effects, in older compared to younger adults. As already observed for arm movements (Poirier et al. 2023a), it is mainly the duration of inactivation that is modulated.

**Figure 8.**
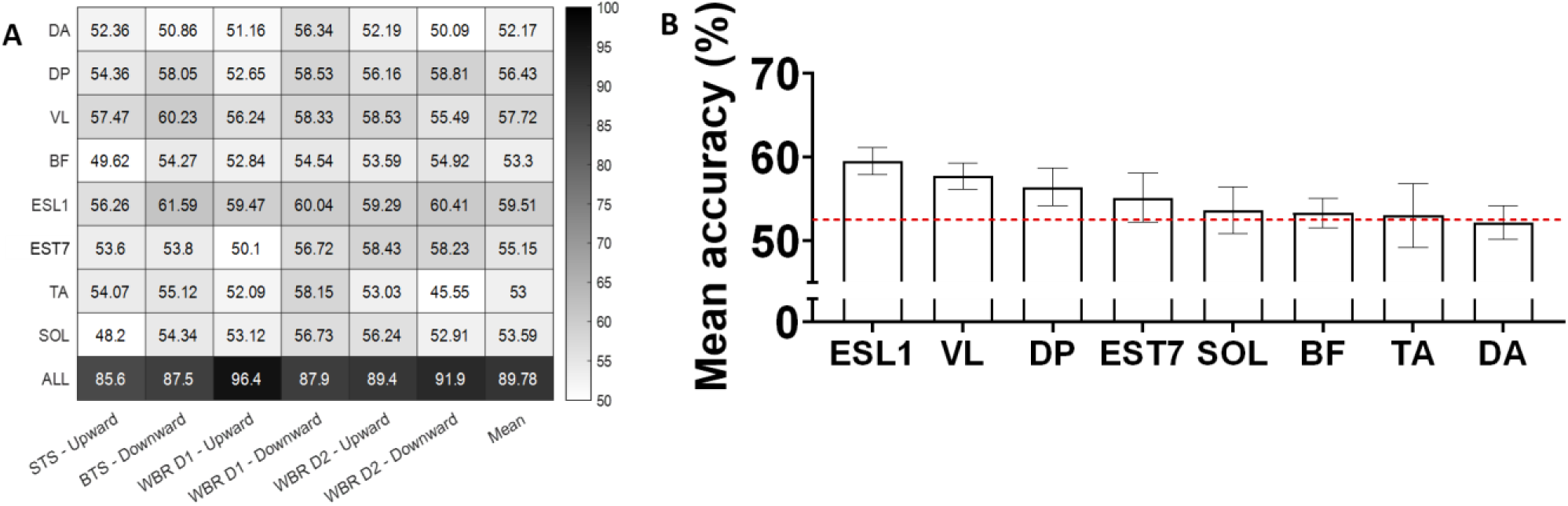
A. Heatmap representing the accuracy of the LDA algorithm to discriminate young from older adults using phasic EMGs recorded during the tasks mobilizing the entire body. The eight first lines correspond to the accuracy of each individual muscle. The last line corresponds to the accuracy of taking all muscles simultaneously (DA: Anterior deltoid, DP: Posterior deltoid, VL: Vastus Lateralis, BF: Biceps Femoris, ESL1: Erector Spinae in L1, EST7: Erector Spinae in T7, TA: Tibialis Anterior, SOL: Soleus). The first six columns correspond to the six whole-body tasks of the experiment (STS = Sit To Stand; BTS = Back To Sit; WBR D1 = Whole Body Reaching near target; WBR D2 = Whole body reaching far target), the last column corresponds to the average accuracy of the six tasks for each muscle. This analysis has been conducted to showcase which muscles are important for discrimination. We can see taht those antigravity muscles (muscles that act against gravity, in our case, the VL and the ESL1) contain relevant information as they reach the highest scores. B. Mean ± SD distance between groups representations by the LDA algorithm for each individual muscle (DA: Anterior deltoid; DP: Posterior deltoid; VL: Vastus Lateralis; BF: Biceps Femoris; ESL1: Erector Spinae in L1; EST7: Erector Spinae in T7; TA: Tibialis Anterior; SOL: Soleus). The higher the distance, the more differentiable the groups are. Values corresponds to the average distance obtain for all six tasks. We can see that those antigravity muscles (muscles that act against gravity, in our case, the VL and the ESL1) contain relevant information as they reach some of the highest values. Error bars correspond to the standard error across a five-fold cross-validation.

**Figure 9.**
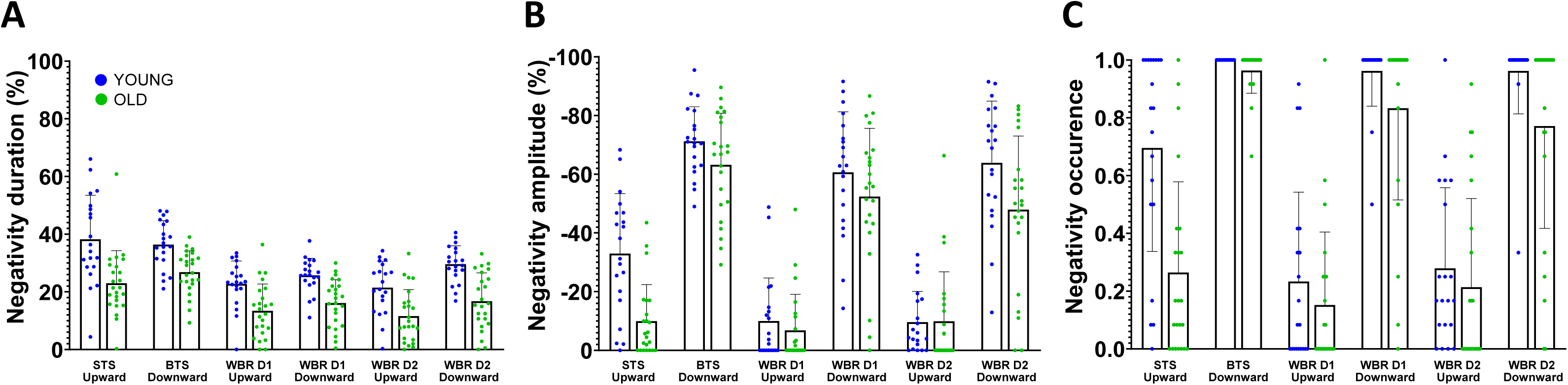
Mean ± SD negativity quantification for all tasks and groups (STS: Seat-to-stand, BTS: Back-to-seat, WBR: Whole-body-reaching, D1: Short distance=15% of the heighy of the subject, and D2: Long distance=30% of the heighy of the subject). Quantification was carried out using three criteria: negativity duration (panel A), negativity amplitude (panel B), and negativity occurrence (panel C) on the antigravity muscles (vastus lateralis and the erector spinae). The blue points correspond to the younger participants, and the green points correspond to the older participants. Each point corresponds to the mean value across trials and antigravity muscles.

Here we were interested in comparing arm movement and whole-body movement control because the scientific literature has reported that the control of whole-body movements changes with age, while the control of arm movements does not (Casteran et al., 2018; Paizis et al., 2008; Poirier et al., 2020, 2023a; Vernazza-Martin et al., 2008). Focusing on a limited number of muscles is problematic, as we risk probing muscles whose activation patterns are not affected by age. To ensure that our restrictive theory-driven analysis provides meaningful results, we verified that our cherry-picked muscles truly conveyed information about age-related modifications of whole-body movement control. To this aim, we employed machine learning analyses that quantified how much each muscle activation was altered by age. This allowed controlling that we were actually focusing on muscles that discriminated movement control between younger and older adults.

Our rationale was the following: if the algorithm can successfully separate the data of younger and older adults, using antigravity muscle activation patterns, this would demonstrate that important information is contained in those muscles regarding age-related modifications of movement control. For more details on similar use and operation of machine learning algorithms on EMG signals, please see (Chambellant et al., 2024; Thomas et al., 2023; Tolambiya et al., 2011). Here we present the results of a Linear Discriminant Analysis (LDA, Johnson & Wichern, 1988) but we verified that we obtained similar conclusions with two other algorithms, namely the Quadratic Discriminant Analysis (QDA, Cover, 1965) and the Support Vector Machine (SVM, Vapnik and Lerner, 1965).

The Machine Learning analysis indeed revealed that antigravity muscles contained important information, allowing separating age-groups with some of the best success-rates (see Figure 8 for results regarding LDA accuracy). The vastus lateralis (VL) and the spinal erectors on L1 (ESL1) achieved the best classification accuracies of 57.72% and 59.51% respectively (considering that these classifications are significantly better than chance if they are above 52.5% according to a fairness test). The main results presented here are therefore quantitatively based. They originate from analyses of the muscles that show the most information to distinguish younger from older adults during whole-body movement. Other muscles, such as DP or EST7, also exhibit reasonably good classification accuracies. This is not unexpected as humans and animals are known to control their varied muscles in a synergistic manner (Berret et al., 2009; d’Avella et al., 2006; Tresch et al., 1999), and even the slightest alteration of movement strategy may require modifying the activation of several muscles.

## Discussion

In younger and older adults, we investigated the muscle activation patterns responsible for arm and whole-body movements. The results revealed an age-related alteration of muscle activation that differed between the types of tasks. Comparing older adults to younger ones, we found that a muscle marker of energetic efficiency was reduced during whole-body movements, but the results show no evidence that this marker was reduced during arm movements. Previous works have demonstrated that this marker allows quantifying the output of a sensorimotor control process that adapts human movements to gravity (Gaveau et al., 2021; Poirier et al., 2022, 2023a). More precisely, this marker allows for quantifying how much one harvests gravity effects to save energy. Here, arm movement results reveal that this energetic-efficiency process remains functional in older adults. During whole-body movements, however, the present results reveal that a criteria linked to energetic-efficiency was downregulated in older adults compared to younger adults. Overall, the present results suggest a compensation process that modulates planning strategies to maximize equilibrium in older adults.

### Age-related compensatory processes in sensorimotor control

A number of studies have proposed that the differences observed between younger and older adults can be interpreted as compensations for age-related deteriorations. Of particular interest are studies from the last decade that have sought to investigate specific motor control processes rather than broad motor performance. For example, some of these studies indirectly suggested that older adults favor feedforward rather than feedback control (Moran et al., 2014; Wolpe et al., 2016) to compensate for the attenuation of sensory processing with increasing age (Moran et al., 2014; Parthasharathy et al., 2022; Saenen et al., 2023). Others indirectly suggested that older adults favor movement efficiency over precision (Healy et al., 2023; Poirier et al., 2020) to compensate for their increased energetic cost (Didier et al., 1993; Hortobagyi et al., 2011; John et al., 2009). Yet, because the focus of these studies was not on compensatory processes, they did not include dedicated experimental conditions. The aim of the present study was to fill this gap.

### Maintained efficiency of arm movements in older adults

The metabolic rate is known to influence resource use, body size, rate of senescence, and survival probability (Brown et al., 2004; DeLong et al., 2010; Strotz et al., 2018; Van Voorhies & Ward, 1999). The nervous system has therefore developed the ability to design movement strategies that minimize our every-day efforts (Cheval et al., 2018; Gaveau et al., 2016; Huang et al., 2012; Morel et al., 2017; Selinger et al., 2015; Shadmehr et al., 2016). The present findings confirm the results of previous arm movement studies that proposed a theory according to which motor control takes advantage of gravity effects to save energy (Berret et al., 2008; Crevecoeur et al., 2009; Gaveau et al., 2014, 2016, 2021; Gaveau & Papaxanthis, 2011). Here, we focused on the muscle activation marker of gravity-related energetic-efficiency, i.e., the negativity of phasic EMG. Previous modeling and experimental work demonstrated that this phasic EMG negativity results from an optimal control process that plans efficient arm movements in the gravity field (Gaveau et al., 2021). As reported by (Poirier et al., 2023a), we found similar phasic EMG negativity during arm movements in older and younger adults. Thus, arm movements equally optimized gravity effects in younger and older adults. These results align with those of studies that probed progressive motor adaptation to a new environment in older adults. Using locally induced force fields in a robotic environment, these studies revealed that older adults decreased their metabolic costs similarly to younger adults while adapting to new environmental dynamics (Healy et al., 2023; Huang & Ahmed, 2014). Overall, results from arm movement studies advocate for the maintenance of the ability to optimally integrate environmental dynamics and plan arm movements that are energetically efficient in older adults.

### Whole-body movements also harvest gravity effects to save energy

Current results also extend the current knowledge on the planning of energetically efficient movements to more global movements, both in younger and older adults. They unravel that deactivating muscles below the tonic level that would be necessary to compensate for external dynamics are not only relevant to controlling focal arm movement but also for whole-body movements. Using a combination of modeling and experimental work, previous studies demonstrated that healthy participants move their arms following trajectories and using muscular patterns that save energy in the gravity environment (Berret et al., 2008; Crevecoeur et al., 2009; Gaveau et al., 2014, 2016, 2021; Gaveau & Papaxanthis, 2011). To isolate gravity effects, most studies focused on one-degree-of-freedom arm movements. Although those studies allowed us to clearly demonstrate how motor planning integrates gravity effects into motor planning, one-degree-of-freedom movements are hardly representative of the rich and complex human movement repertoire. The present study, using more ecological movements, basically extends the optimal integration of gravity effects theory to every-day movements.

### Decreased efficiency of whole-body movements in older adults

Contrary to focal arm movements, we observed a strong age difference during global movements that engaged the entire body, here sit to stand / back to sit and whole-body reaching movements. Specifically, the negativity of phasic EMG was significantly reduced in older compared to younger adults. This suggests that whole-body movements are less energetically efficient in older adults than in younger ones, adding to the general result that global movements are more energy-demanding for older adults compared to younger adults (Didier et al., 1993; Hortobagyi et al., 2003, 2011; John et al., 2009). Previous kinematic studies suggested that older adults favor movement strategies that maximize balance maintenance rather than energy efficiency (Casteran et al., 2018; Paizis et al., 2008). However, age differences observed during whole-body movements may also be interpreted as an inability to save energy when coordinating complex movements (Goodpaster et al., 2006; Henry & Baudry, 2019; Quinlan et al., 2018; Vernazza-Martin et al., 2008). Here, contrasting results from arm and whole-body movements in the same participants, we provide support for a compensation process that adapts movement strategy in older adults, rather than a deterioration of the ability to optimally coordinate whole-body movements. Since arm movements revealed that older participants maintained the ability to plan energetically efficient movements, altered whole-body movement may be explained as an adaptation of movement strategy rather than deteriorated motor planning. More importantly, we found that decreased efficiency was associated with decreased center-of-mass displacement and speed, i.e., less instability. This further suggests that decreased efficiency in older adults is a compensation process that trades efficiency with equilibrium maintenance. This could be explained as an optimal motor planning process that minimizes a composite cost function; i.e., energy and unstability. It has been proposed that the central nervous system combines different costs – related to energy, precision, or duration, for example – when planning a movement (Berret et al., 2011; Gielen, 2009; Healy et al., 2023; Liu & Todorov, 2007; Mombaur et al., 2010; Poirier et al., 2023a; Tanis et al., 2023; Vu et al., 2016). In older adults, this combination would increase the relative weighting of the instability (equilibrium) cost and decrease the relative weighting of the energetic cost. Future work may use this framework to probe age-related motor adaptation.

### Effect of target distance

During the whole-body reaching task, reusing the protocol (Casteran et al., 2018), we varied the antero-posterior distance of the target to be reached. Casteran et al. (2018) found larger differences between younger and older participants when the target was further. Consequently, we hypothesized that the further away the target, the greater the age differences in the negativity epochs of phasic EMGs. The present results do not validate this hypothesis (see Supplementary Table 3).

### Age-related compensation in the brain

In the sensorimotor field, following the consensus that aging is associated with increased activation and increased spatial recruitment, numerous studies have attempted to establish a correlation between brain activation and behavioral performance in older adults (for reviews, see Fettrow et al., 2021; Poirier et al., 2021; Seidler et al., 2010; Ward, 2006). This literature has not reached a consensus on the neural changes underlying compensatory mechanisms in the aging brain. Several studies reported a positive correlation (Cassady, Gagnon, et al., 2020; Clark et al., 2014; Harada et al., 2009; Heuninckx et al., 2008; Holtzer et al., 2015; Jor’dan et al., 2017; Larivière et al., 2019; Mattay et al., 2002; Spedden et al., 2019), and as many reported no correlation or even a negative correlation (Bernard & Seidler, 2012; Cassady et al., 2019; Cassady, Ruitenberg, et al., 2020; Fernandez et al., 2019; Hawkins et al., 2018; Holtzer et al., 2016; Loibl et al., 2011; Riecker et al., 2006; Ward et al., 2008). Building on the theoretical work of Krakauer et al. (2017), we recently proposed that an important reason for this lack of consensus may be that previous studies, while focusing on brain activations, used crude behavioral paradigms that likely mixed deteriorated and compensatory processes (Poirier et al., 2021). Using behavioral paradigms that focus on specific motor control processes, as performed here, could help differentiate compensatory mechanisms from deteriorative ones.

### Role of physical and cognitive fitness in age-related compensation

Physical and cognitive fitness may influence how much older adults favor stability over energetic efficiency. It is well-known that physical and cognitive fitness significantly impact functional mobility in older adults (Marusic et al., 2018; Wickramarachchi et al., 2023; Zhao et al., 2014). One could speculate that physical and cognitive fitness are inversely related to the level of physical and cognitive deterioration. For example, muscle force and sensory integration are crucial for controlling balance. The more deteriorated they are, the greater the need for compensatory processes to adapt movement control to the participant’s capacities. Future research should account for variations in physical and cognitive fitness to better understand their role in the development of compensatory mechanisms.

### Simple mono-articular vs complex multi-articular arm movements

Another aspect that needs to be highlighted here is the choice of the arm task, which is not representative of all existing tasks for studying upper limb motor skills. Using this very same task, the results from two previous studies also support the preservation of arm movement efficiency in older adults (Poirier et al., 2020, 2023a). One may wonder whether the present conclusions would hold for more complex arm movements. Using multi-degree of freedom arm movements to study motor adaptation to an externally imposed force-field, other studies also reported results showing that, alike younger adults, older adults maintain the ability to produce movements that are energetically efficient (Healy et al., 2023; Summerside et al., 2024). The present mono-articular results are therefore likely to generalize to other types of arm movements. In young adults, the efficient integration of gravity effects to save energy has been demonstrated with varied arm movements, such as single or muti-degree of freedom pointing movements, drawing movements, reach to grasp movements, or arm movements that transport a hand-grasped object (Berret et al., 2008; Crevecoeur et al., 2009; Gaveau et al., 2011; Gaveau & Papaxanthis, 2011; Le Seac’h & McIntyre, 2007; Papaxanthis et al., 1998, 2005; Yamamoto & Kushiro, 2014b). Future work may test whether the present conclusions extend to more complex and functional arm movements.

In conclusion, probing a specific motor control process, the present study provides a set of behavioral results that support the interpretation of a compensatory process that counterbalances other deteriorated processes in older adults. Probing age effects on specific sensorimotor control processes may help disentangle compensation from deterioration processes that occur through healthy aging (Poirier et al., 2021). We believe that understanding compensation at a behavioral level is an important step toward pinpointing its neural underpinning (Krakauer et al., 2017) and, later, preventing unhealthy aging (Baltes & Baltes, 1990; Martin et al., 2015; Zhang & Radhakrishnan, 2018).

## Acknowledgements

We thank Yves Ballay, Denis Barbusse, and Gabriel Poirier for their support during the pilot study. We also thank all the participants who took part in the experiment.

## Data, scripts, code, and supplementary information availability

Data are available online: 10.5281/zenodo.10619701, webpage hosting the data: https://doi.org/10.5281/zenodo.10619701 (*citation of the data eg* Mathieu et al, 2024);

Scripts and code are available online: 10.5281/zenodo.10634004, webpage hosting the scripts: https://doi.org/10.5281/zenodo.10634004 (*citation of the scripts eg* Mathieu et al, 2024);

Supplementary information is available online: 10.5281/zenodo.10671496, webpage hosting the file: https://doi.org/10.5281/zenodo.12671953 (*citation of the supplementary file eg* Mathieu et al, 2024);

## Conflict of interest disclosure

The authors declare that they comply with the PCI rule of having no financial conflicts of interest in relation to the content of the article.

Jérémie Gaveau is a member of the managing board of the PCI Health & Movement Sciences.

