## Supplemental for "Comparing arm to whole-body motor control disambiguates age-related deterioration from compensation"

### Supplementals

| Subjects | | ARM | STS/BTS | | | | WBR D1 | | | | WBR D2 | | | |
|---|---|---|---|---|---|---|---|---|---|---|---|---|---|---|
| | | DA | VL | | ESL1 | | VL | | ESL1 | | VL | | ESL1 | |
| | | Right | Right | Left | Right | Left | Right | Left | Right | Left | Right | Left | Right | Left |
| Young | 1 | X | X | X | X | NO | X | X | X | NO | X | X | X | NO |
| | 2 | X | X | X | X | X | X | X | Slow | Slow | X | X | X | X |
| | 3 | X | X | X | X | NO | X | X | X | NO | X | X | X | NO |
| | 4 | X | X | X | X | NO | X | X | X | NO | X | X | X | NO |
| | 5 | X | X | X | X | NO | X | X | X | NO | X | X | X | NO |
| | 6 | X | X | NO | NO | NO | X | X | X | NO | X | X | X | NO |
| | 7 | X | X | X | X | NO | X | X | X | NO | X | Slow | Slow | NO |
| | 8 | X | X | X | X | X | X | X | X | X | X | X | X | X |
| | 9 | X | X | X | X | X | X | Slow | X | Slow | X | X | X | X |
| | 10 | X | X | X | X | X | X | X | X | X | X | X | X | X |
| | 11 | X | X | X | X | X | X | X | X | X | X | X | X | X |
| | 12 | X | X | X | X | X | X | X | Slow | Slow | X | X | X | X |
| | 13 | X | X | NO | X | X | Fast | NO | X | X | X | X | X | X |
| | 14 | X | X | X | X | X | X | X | X | X | X | X | X | X |
| | 15 | X | X | X | X | X | X | X | X | X | X | X | X | X |
| | 16 | X | X | X | X | Slow | X | X | X | X | X | X | X | X |
| | 17 | X | X | X | X | X | X | X | NO | NO | X | X | X | X |
| | 18 | X | X | X | X | X | X | X | Slow | Slow | X | X | Slow | Slow |
| | 19 | X | X | X | X | X | X | X | X | X | X | X | X | X |
| | 20 | X | X | X | X | X | X | X | X | X | X | X | X | X |
| Old | 1 | X | X | X | X | Fast | X | X | X | X | X | Fast | X | X |
| | 2 | X | X | X | X | X | X | X | X | X | X | X | X | X |
| | 3 | X | X | X | X | X | X | X | X | X | X | X | X | X |
| | 4 | X | X | X | X | X | X | X | X | NO | X | Slow | Slow | NO |
| | 5 | X | X | X | X | X | NO | Slow | Slow | Slow | NO | X | X | X |
| | 6 | X | X | X | X | X | Slow | X | NO | NO | X | X | NO | NO |
| | 7 | X | X | X | X | X | X | X | Slow | X | X | X | X | X |
| | 8 | X | X | X | X | X | X | X | X | X | X | X | X | X |
| | 9 | X | X | NO | X | X | Fast | NO | X | X | Fast | NO | X | X |
| | 10 | X | X | X | X | X | X | NO | X | X | X | NO | X | X |
| | 11 | X | NO | NO | X | X | NO | NO | X | X | NO | NO | X | X |
| | 12 | X | Slow | X | NO | X | X | X | NO | X | X | X | NO | X |
| | 13 | X | X | X | X | X | X | NO | X | X | Fast | NO | X | X |
| | 14 | X | X | X | X | X | X | X | X | X | X | X | X | X |
| | 15 | X | X | X | X | X | X | X | X | X | X | X | X | X |
| | 16 | X | X | X | X | X | X | X | X | X | X | X | X | X |
| | 17 | X | X | X | X | X | X | X | X | X | X | X | X | X |
| | 18 | X | X | X | Slow | X | X | NO | X | X | X | X | NO | NO |
| | 19 | X | X | X | X | X | X | X | X | X | Fast | Fast | Fast | Fast |
| | 20 | X | X | X | X | X | X | X | X | X | X | X | X | X |
| | 21 | X | X | X | X | X | X | X | X | X | X | X | X | X |
| | 22 | X | X | X | X | X | X | X | X | X | X | X | X | X |
| | 23 | X | X | X | X | X | Slow | Slow | Slow | Slow | X | X | X | X |
| | 24 | X | X | X | X | X | X | X | X | X | X | X | X | X |

**Supplementary Table 1.** Some of the EMG recordings exhibited aberrant values. This table reports the quality of EMG recordings for the main antigravity muscles in each participant (numbered on the left), and each task, i.e. DA (Deltoid Anterior head) during focal arm movements, and VL (Vastus Lateralis) and ESL1 muscles (Erector Spinae at Lumbar 1 level) during global whole-body movements. "X": all trials were correct. "Fast": some of the fast trials, maximum 4/12, have been excluded. "Slow": some of the slow trials, maximum 2/6, have been excluded. "NO": too many trials exhibited aberrant values and, therefore, the muscle was not used for the analyses presented in this manuscript. STS/BTS: Seat-to-stand/Back-to-seat. WBR D1: Whole body reaching near target. WBR D2: whole body reaching far target.

| Task | | Direction | Group | Mean (s) ± SD (s) |
|---|---|---|---|---|
| ARM | Tonic | Upward | OLD | 6,104± 2,8 |
| | | | YOUNG | 3,907± 1,4 |
| | | Downward | OLD | 6,395± 3,6 |
| | | | YOUNG | 3,544± 1,6 |
| | Fast | Upward | OLD | 0,221± 0,08 |
| | | | YOUNG | 0,210± 0,08 |
| | | Downward | OLD | 0,249± 0,10 |
| | | | YOUNG | 0,213± 0,08 |
| STS/BTS | Tonic | Upward | OLD | 2,144± 0,7 |
| | | | YOUNG | 1,698± 0,5 |
| | | Downward | OLD | 2,858± 1,0 |
| | | | YOUNG | 2,141± 0,6 |
| | Fast | Upward | OLD | 0,357± 0,09 |
| | | | YOUNG | 0,267± 0,05 |
| | | Downward | OLD | 0,396± 0,11 |
| | | | YOUNG | 0,279± 0,04 |
| WBR D1 | Tonic | Upward | OLD | 4,009± 1,2 |
| | | | YOUNG | 3,106± 1,0 |
| | | Downward | OLD | 3,375± 0,9 |
| | | | YOUNG | 2,739± 0,7 |
| | Fast | Upward | OLD | 0,521± 0,11 |
| | | | YOUNG | 0,419± 0,05 |
| | | Downward | OLD | 0,458± 0,07 |
| | | | YOUNG | 0,389± 0,05 |
| WBR D2 | Tonic | Upward | OLD | 4,083± 1,3 |
| | | | YOUNG | 3,276± 0,9 |
| | | Downward | OLD | 3,594± 1,2 |
| | | | YOUNG | 2,747± 0,8 |
| | Fast | Upward | OLD | 0,529± 0,10 |
| | | | YOUNG | 0,445± 0,06 |
| | | Downward | OLD | 0,525± 0,09 |
| | | | YOUNG | 0,442± 0,05 |

**Supplementary Table 2.** Mean movement durations ± (SD) are presented for each task, movement direction, movement speed and age-group. STS/BTS = Sit To Stand / Back To Sit; WBR D1 = Whole-Body Reaching near target; WBR D2 = Whole-Body reaching far target.

| | Effects | F | p-value | Effect size (partial $\eta^2$ ) |
|---|---|---|---|---|
| **A. Movement duration** | Type of Tasks effect | 433 | 9,38E-24 | 0,911 |
| | Age effect | 14,5 | 4,58E-04 | 0,256 |
| **ANCOVA Age x Type of Tasks** | Age x Type of Tasks effect | 20,0 | 5,72E-05 | 0,322 |
| **B. Negativity Index** | Type of Tasks effect | 0,53 | 4,70E-01 | 0,013 |
| | Age effect | 0,54 | 4,68E-01 | 0,013 |
| **ANCOVA Age x Whole-body tasks** | Age x Type of Tasks effect | 5,48 | 2,44E-02 | 0,120 |
| **C. Negativity Index** | Whole-body Tasks effect | 0,78 | 9,30E-01 | 0,002 |
| | Age effect | 4,50 | 4,00E-02 | 0,103 |
| **ANCOVA Age x Type of Tasks** | Age x Whole-body Tasks effect | 0,77 | 4,67E-01 | 0,019 |
| **D. Negativity Duration** | Whole-body Tasks effect | 0,32 | 7,20E-01 | 0,009 |
| | Age effect | 21,5 | 4,54E-05 | 0,374 |
| **ANCOVA Age x Whole-body tasks** | Age x Distance effect | 2,49 | 8,99E-02 | 0,065 |
| **E. Negativity Amplitude** | Whole-body Tasks effect | 0,63 | 5,37E-01 | 0,017 |
| | Age effect | 1,16 | 2,80E-01 | 0,031 |
| **ANCOVA Age x Whole-body tasks** | Age x Distance effect | 0,72 | 4,89E-01 | 0,020 |
| **F. Negativity Frequency** | Whole-body Tasks effect | 0,39 | 6,75E-01 | 0,011 |
| | Age effect | 3,62 | 6,50E-02 | 0,091 |
| **ANCOVA Age x Whole-body tasks** | Age x Distance effect | 0,358 | 7,01E-01 | 0,010 |

**Supplementary Table 3.** Details of the statistical analyses presented in the manuscript for **A**. the movement duration, **B**. the negativity index for whole-body tasks, **C**. the negativity index for the comparison arm and whole-body tasks, **D**. the negativity duration for whole-body tasks, **E**. the negativity amplitude for whole-body tasks, and **F**. the negativity frequency for whole-body tasks. Are gathered in the table the F, the p-value and the effect size (partial $\eta^2$).

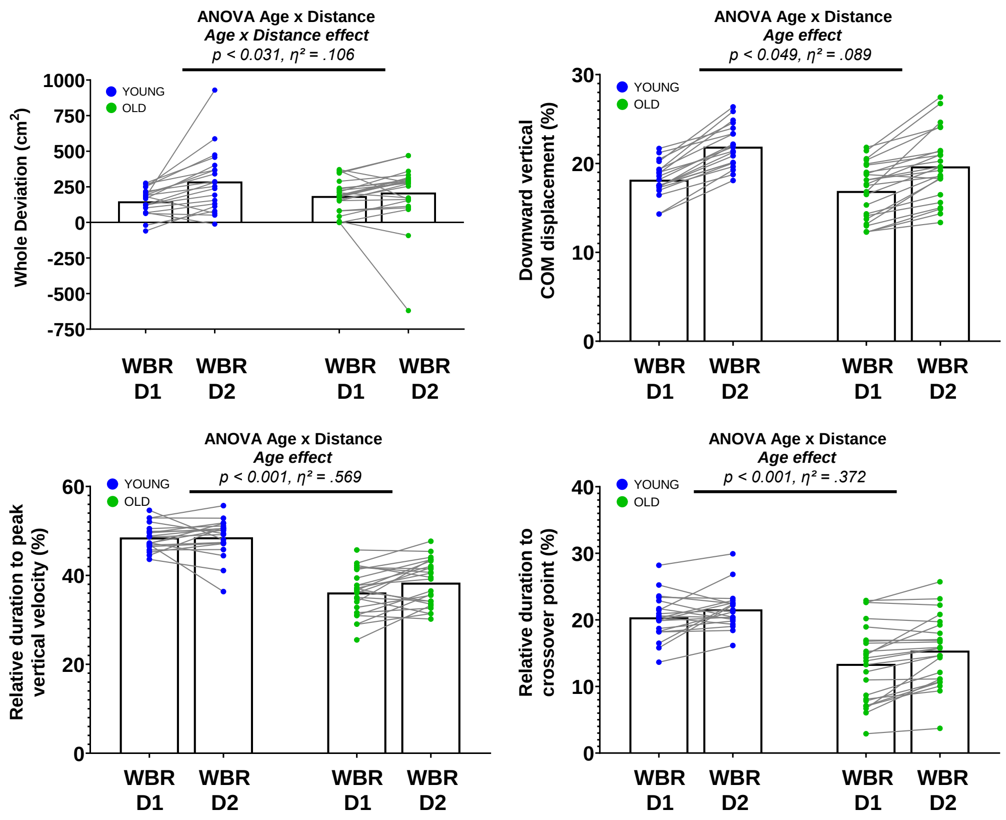


**Supplementary Figure 1.** Reproduction of the analyses of Paizis et al. (2008) (panels **A** and **B**) and Casteran et al. (2018) (panels **C** and **D**). Both studies showed age-related differences in motor skills during balance tasks (downward whole-body reaching tasks).

**Paizis et al. (2008)**. We computed the two main criteria used by Paizis et al. (2008): the whole deviation and the downward vertical center of mass (CoM) displacement. The whole deviation was computed using a trapezoidal numerical integration from Matlab [Mat, 2021] to obtain the area between the actual trajectory from the wrist and the straight path between movement start and end. The downward vertical displacement was computed on the CoM using the mathematical seven-segment model of the body detailed in the manuscript between movement start and end and was normalized by the height of the participant. Paizis et al. (2008) only tested one distance. We observe an interaction Age x Distance effect where Paizis et al. (2008), with only one distance, only observe an Age effect. We observe the same Age effect as they did for the vertical CoM displacement, with the younger participants having more displacement than the older ones. The results we found are in direct agreement with those obtained by the authors.

**Casteran et al. (2018)**. We computed the two main criteria used by Casteran et al. (2018): the relative duration to peak vertical velocity of the CoM and the relative duration to crossover point. The relative duration to peak vertical velocity of the CoM was obtained using the mathematical seven-segment model of the body detailed in the manuscript between movement start and end. The crossover point between the anteroposterior and vertical velocities was obtained by decomposing the CoM velocity profile into its Antero-Posterior (ΔAP trajectory) and Vertical (ΔV) components. The point of interest is the moment when the vertical component surpasses the antero-posterior one. Casteran et al. (2018) tested the same two distances. We observe an Age effect for the relative time to peak vertical velocity, as Casteran et al. (2018) did, suggesting a vertical peak velocity occurring earlier for older participants. We found an Age effect for the crossover point between vertical and antero-posterior components of the CoM velocity, while Casteran et al. (2018) suggested that – but did not directly test – there may exist an interaction effect Age x Distances. The present results do not validate the supposed interaction effect. Distance does not seem to more strongly affect the movements of older adults compared to younger adults.
